# The Pannexin-1 N-terminal Helix Gates a Switch between Ion Conductance and Anandamide Transport

**DOI:** 10.1101/2024.03.11.584466

**Authors:** Connor L. Anderson, Nicolas L. Weilinger, Frank Visser, Allison C. Nielsen, Andrew K.J. Boyce, Roger J. Thompson

**Affiliations:** Department of Cell Biology and Anatomy, Hotchkiss Brain Institute, Cumming School of Medicine, University of Calgary

## Abstract

Anandamide is an endovanilloid and endocannabinoid with ligand activity at transient receptor potential vanilloid 1 channels and cannabinoid receptors, respectively. We have reported that block of Pannexin-1 channels in the CA1 hippocampus can increase concentrations of anandamide and induce presynaptic plasticity. It is not known how an ion channel can contribute to clearance of a lipid-derived signalling molecule. Here, we use electrophysiology and imaging of uptake of fluorescent anandamide to determine the structure-function relationship between the ion conduction and anandamide trasporter activities of pannexin-1. Expression of rat, mouse or human pannexin-1 in HEK cells caused a time dependent increase in anandamide uptake by all three orthologs. However, human pannexin-1 had reduced ion conduction. Low concentrations of anandamide augmented uptake of its fluorescent derivative, whereas higher concentrations competed, suggesting that anadamide may facilitate its own transport. Deletion of the N-terminal helix of pannexin-1 and the channel blocker, probenecid, blocked ion conduction but enhanced anandamide transport. In contrast, mutation of pore facing isoleucine 41 caused a gain of function in ion conduction with loss of anandamide transport. We conclude that the pannexin-1 channel is a dual ion channel / anandamide transporter and that these properties are gated by the channels N-terminal helix and likely linked to its presence or absence within the pore lining region.

## Introduction

The ion / small molecule permeable channels formed by pannexin-1 (Panx1) have roles in cell death^1,2^, purinergic signalling by ATP release^3,4^, and endovanilloid signalling by controlling anandamide (AEA) concnetrations in the CA1 hippocampus^5^. Panx1 is expressed at the postsynaptic density (PSD) of hippocampal and cortical pyramidal neurons^6^, where it can enhance high-frequency stimulation-induced long-term potentiation (LTP) in the hippocampus and attenuate low-frequency induced long-term depression (LTD)^7^. We reported a role for postsynaptic Panx1 in activity-dependent glutamate release^5^. In this model, Panx1 controls extracellular concentrations of AEA, which increased during Panx1 block and activated presynaptic transient receptor potential vanilloid 1 (TRPV1), leading to seconds-long high-frequency excitatory postsynaptic currents. However, the mechanism of how Panx1 contributes to clearance of hydrophobic AEA was not identified. Understanding this mechanism could lead to discovery of a novel endocannabinoid / endovanilloid transporter and present new opportunities to pharmacologically manipulate AEA signalling.

The transport of AEA across membranes has been extensively studied and several mechanisms have been proposed^8^. These include direct diffusion through the plasma membrane and facilitated transport by proteins. One such example is an enzymatically inactive isoform of fatty acid amide hydrolase (FAAH), called FLAT^9^. There is evidence for direct diffusion through the membrane being facilitated by cholesterol^10^. In most cases of the proposed transport mechanisms, AEA metabolism by FAAH or binding to intracellular fatty acid binding proteins helps to maintain an intracellular ‘sink’ for a sustained inward driving force^11–14^. Diffusion through the plasma membrane would not be amenable to regulation that is required for termination of synaptic signals and supports the idea of specific endocannabinoid membrane transporters (EMT), which have not been identified.

Two key properties of the unidentified EMT are suggestive of a channel-like mechanism. The first is a low temperature dependence for AEA uptake; the Q_10_ for AEA transport is 1.4^15,16^, which is in the range for most ion channels including other ATP permeable channels^17^. The second is that AEA uptake is widely believed to be energy independent because it occurs without ATP hydrolysis and ionic gradients are not required^15^, suggesting a facilitated diffusion-based mechanism. Based on these observations, taken with our report that Panx1 blockers increase AEA concentrations in hippocampal slices^5^, we predicted that Panx1 directly transports AEA. Recently published cryo-EM structures of Panx1 were used to guide mutagenesis of the channel and assess function^18,19^, leading us to propose a model where Panx1 switches between ion conductance and AEA transport. The two conductance modes are gated by the N-terminal domain and a key intrapore isoleucine residue, I41. We further propose, based on the closed channel configuration reported by Kuzuya et al, that the AEA permeable / Cl^-^ impermeable state may arise when membrane lipids have migrated into the pore to ‘close’ the channel.

## Results

### Pannexin-1 orthologs enhance uptake of fluorescent anandamide

To determine if Panx1 contributes to AEA transport across the plasma membrane, we transfected HEK 293T cells with rat (r), mouse (m) or human (h) Panx1 and utilized CAY10455, an AEA molecule conjugated to fluorescein isothiocyanate (FITC)^20^. This was necessary to assess putative AEA transport because AEA (and CAY10455) is non-charged, preventing the measurement of flux with electrophysiological techniques. CAY10455 is non-fluorescent until cytosolic esterases cleave the AEA-FITC linker, allowing FITC to become fluorescent and quantified as a surrogate for AEA transport. Panx1 cDNA was co-transfected with the fluorescent reporter mKate, to identify positively transfected cells for experiments. To isolate Panx1 function and confirm membrane expression, we performed whole-cell patch-clamp (− 80mV to +80mV voltage ramps) to record voltage-sensitive currents and membrane biotinylation to quantify the relative membrane expression of the constructs. Carbenoxolone (CBX; 100 µM) was used to confirm currents were due to Panx1. All three Panx1 orthologs were detected by membrane biotinylation, with low levels of endogenous hPanx1 in non-transfected HEK 293T cells (**Supplementary Fig 1**). In these non-transfected control cells, minimal CBX-sensitive currents were recorded (**Fig. 1AB**). Weakly CBX sensitive currents were also recorded from HEK 293T cells expressing hPanx1 (**Fig 1AB**), consistent with previous reports that hPanx1 has weak ion channel activity^4,21,22^. Recordings from cells expressing rPanx1 or mPanx1 showed robust CBX-sensitive voltage-activated currents (**Fig 1AB**).

**Fig. 1.**
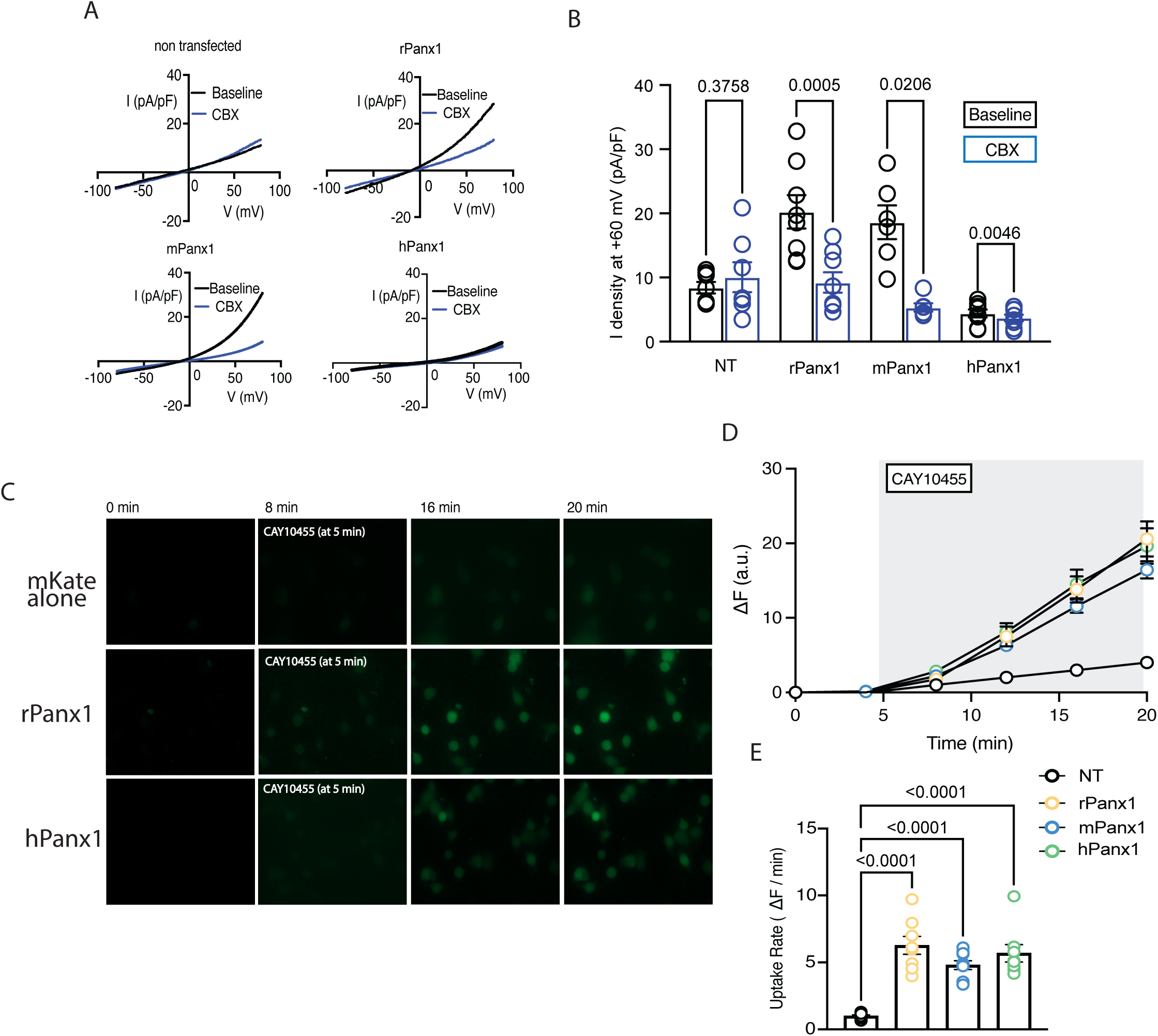
Panx1 enhances uptake of fluorescent anandamide. Whole-cell voltage ramp recordings from GFP positive HEK 293T cells transfected with rat, mouse or human (rPanx1, mPanx1, hPanx1) orthologs. The rodent orthologs show pronounced carbenoxolone (CBX; 100 μM) sensitive currents, indicating Panx1 expression. **b.** Quantification of whole-cell current density at +60 mV for each Panx1 ortholog. Non-transfected (NT) cells had no significant carbenoxolone (CBX; 100μM) sensitive current (two-tailed paired t-test, n=7, P=0.3758, t=0.9565). Significant (two tailed paired t-test) CBX-sensitive currents were seen in rat (n=8, P=0.0005, t=6), mouse (n=6, P=0.0206, t=3.352) and human (n=8, P=0.0046, t=4.091) Panx1 expressing HEK 293T cells. **c.** Sample images of HEK 293T cells during uptake of 500 nM CAY10455. **d**. Time course of uptake of CAY10455 for each Panx1 ortholog. Number of experimental replicates are, for NT n=6; rPanx1 n=8; mPanx1 n=10; hPanx1 n=7. **e**. comparison of the rate of CAY10455 uptake for each ortholog of Panx1 to NT control cells. Significant increases (one way ANOVA, n’s as in d. P<0.0001, F=23.17, df=35) in uptake are seen for rPanx 1, mPanx1 and hPanx1 (P value vs. NT are shown in the plot), with no differences between the Panx1 orthologs.

Having confirmed that all Panx1 orthologs traffic to the membrane and form functional channels, we next used CAY10455 uptake and subsequent fluorescence as a proxy for AEA uptake. After a 5 min baseline, 500 nM CAY10455 was added to the bath and imaged for 15 minutes to quantify uptake (**Fig. 1C**). All mKate positive cells in the field of view were averaged as a single experimental replicate and CAY10455 fluorescence intensity for all replicates was averaged at each time point (**Fig 1D**), then quantified as the rate of uptake (i.e. slope; ΔF/min;) from 8 to 20 min (**Fig 1E**). All Panx1 orthologs, including hPanx1, displayed robust increases in fluorescence compared to cells transfected with mKate alone (**Fig. 1DE**), suggesting CAY10455 influx and thus, AEA transport, is facilitated by Panx1 expression.

It was reported that CAY10455 and AEA compete for transport by the unidentified EMT^20^. Therefore, we tested if co-application of AEA reduced CAY10455 uptake by applying increasing concentrations of AEA in the presence of 500nM CAY10455. Interestingly, low concentrations (1 and 10 nM) significantly increased the rate of CAY10455 uptake (**Fig 2AB**), whereas 100nM AEA inhibited CAY10455 uptake to a level like that seen in non-transfected cells (**Fig 2AB**). This indicates that AEA may facilitate its own uptake at low concentrations and compete with CAY10455 uptake at 100 nM (**Fig. 2AB**). We evaluated if 100 nM AEA affected whole-cell currents carried by Panx1. In HEK 293T cells expressing rPanx1, 100nM AEA did not alter Panx1 current (**Fig. 2CD)**, suggesting that ion currents and AEA transport may be functions of Panx1 that arise from distinct conformational states.

**Fig 2.**
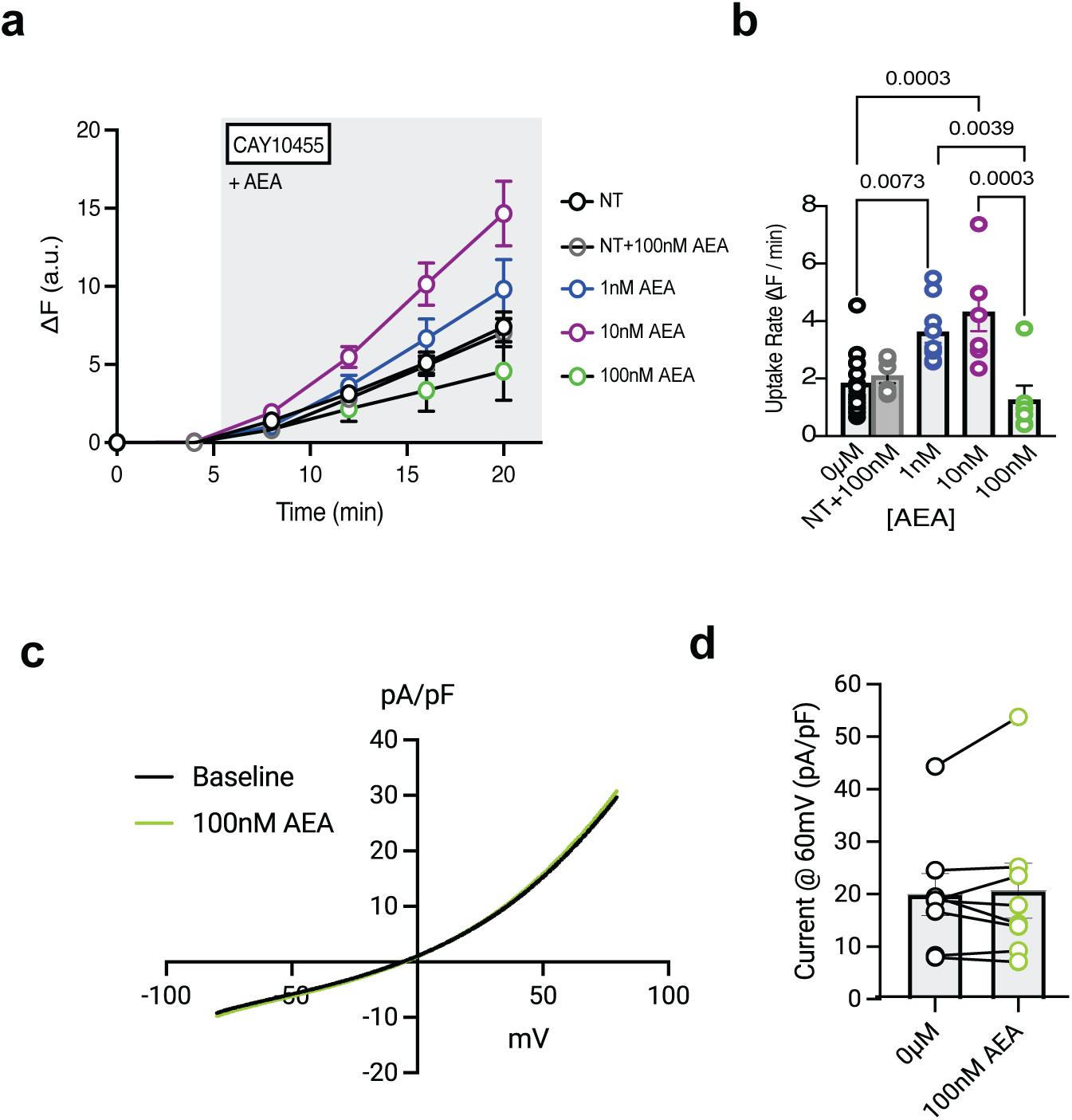
Anandamide competes with CAY10455 for uptake into Panx1 expressing HEK 293T cells. **a**. Plots of the time course of CAY10455 uptake in the presence of 3 different concentrations of anandamide (AEA). Low nM concentrations facilitated CAY10455 uptake while 100 nM inhibited it. Experimental replicates (i.e. fields of view) are NT=12, NT+100 nM AEA=6, 1 nM AEA=9, 10 nM AEA=7, 100 nM AEA=6. Error bars are SEM. **b**. Rates of CAY10455 uptake, calculated as the slope of the plots in a, in presence of increasing doses of AEA. Comparison was made with one-ANOVA (n’s as in a. P<0.0001, F=9.584, df=42) and only significant comparisons are indicated as P values in the plot. **c**. whole cell recordings of HEK 292T cell expressing rPanx1 in the absence or presence of 100 nM AEA. **d**. current density of whole cell recordings in the absence and presence of AEA. No effect of AEA on rPanx1 ion currents is seen (student’s t test, n=8, P=0.3970, t=0.9021).

### Differential effects of Panx1 blockers on ionic currents and AEA uptake

Two conventional Panx1 blockers, probenecid (PBN) and carbenoxolone (CBX) block ion and adenosine triphosphate (ATP) conduction^23–25^. However, we reported that PBN augments the uptake of ethidium bromide (EtBr) by Panx1-expressing cells, yet CBX had no effect^26^, suggesting that blockers of Panx1 may differentially affect ion and small molecule transport by the channels. Here we evaluated if PBN and CBX affected Panx1’s AEA transport. CBX blocked ion currents (**Fig 1AB**) but like conduction of EtBr, CBX did not alter uptake of CAY10455 (**Fig 3A**). In contrast, PBN blocked ion currents (**Fig 3B**) and significantly increased CAY10455 uptake (**Fig. 3CD**). This suggests that in the presence of ion channel block by PBN, but not CBX, Panx1 is in a conformational state that favours AEA transport.

**Fig 3.**
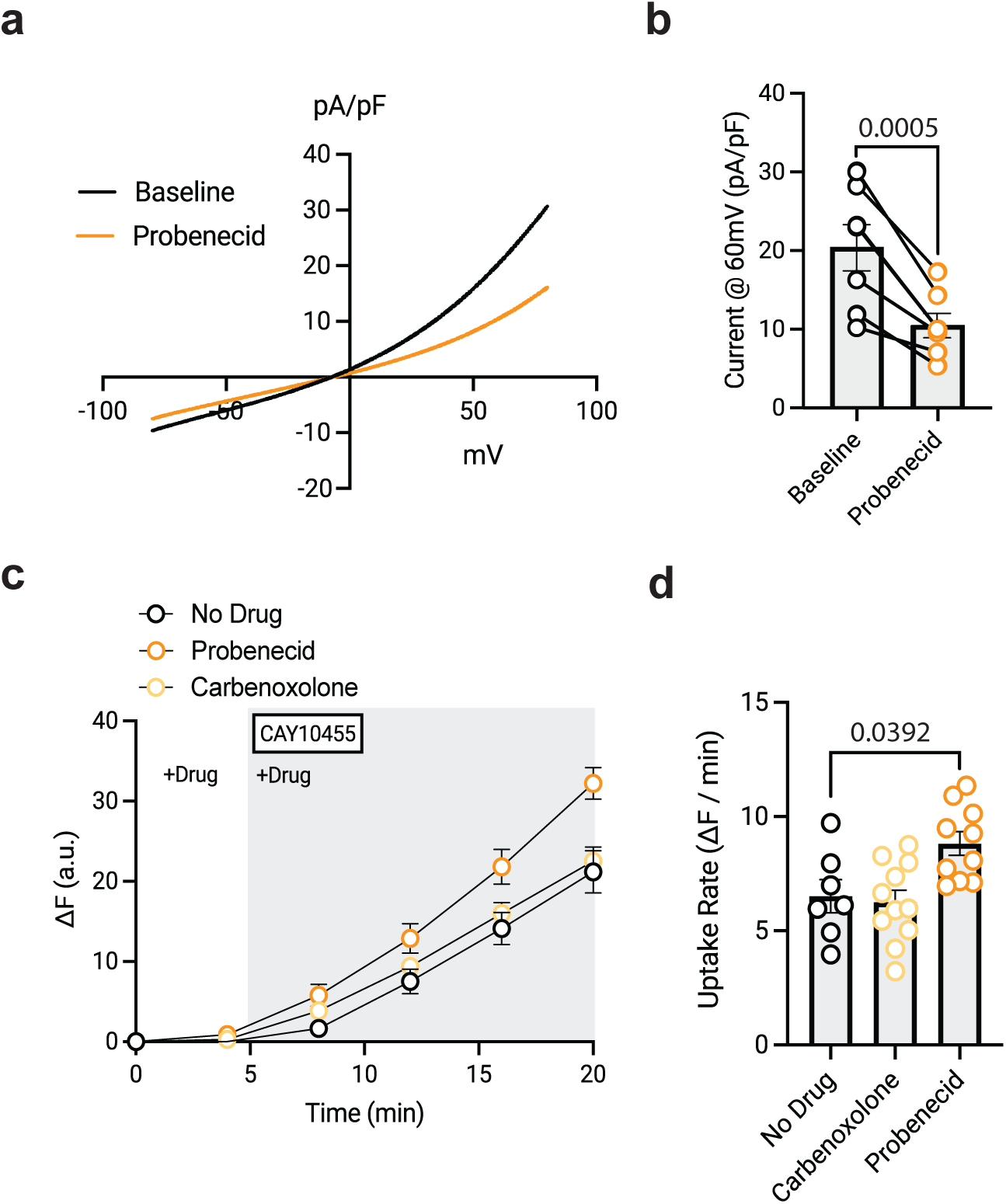
Different effects of Panx1 blockers on CAY10455 uptake. **a.** Similar to carbenoxolone (see Fig 1), the blocker probenecid (1 mM) inhibits whole cell currents carried by Panx1. **b**. quantification of the effects of probenecid on current density in rPanx1 expressing HEK 292T cells. Two-tailed student’s t test, n=8, P=0.0005, t=6.152). **c.** time course plots of the effects of the Panx1 blockers, carbenoxolone and probenecid on CAY10455 uptake by rPanx1. Number of replicates were, no drug n=7, carbenoxolone n=11, probenecid n=10. **d**. rates of CAY10455 uptake by rPanx1 in the presence of either carbenoxolone (100 μM) or probenecid (1 mM). Note that carbenoxolone does not alter CAY10455 uptake and probenecid enhances it. Comparisons were by one-way ANOVA (n’s as in a. P<0.0056, F=6.418, df=27). Direct comparisons, no drug vs carbenoxolone (P>0.9999) and no drug vs probenecid (P=0.0392).

### The Panx1 N-terminal helix gates a switch between ion conduction and AEA transport

Cryo-EM structures of Panx1 resolved in the presence of CBX or PBN indicate that the two blockers likely have distinct mechanisms of action^18,19^. For example, Kuzuya and colleagues reported that PBN closes Panx1 to ion conduction by a conformational rearrangement of the N-terminal helix (NTH), which normally resides in the pore but moves into the cytosol during channel closure^19,27^. Interestingly, Kuzuya et al also suggested that membrane lipids move between Panx1 subunits to fill the pore region and prevent ion conductance. In contrast, Ruan et al reported that CBX binds to the extracellular facing selectivity filter at W74 and likely acts as a pore blocker^18^. A conformational change in the Panx1 NTH in the presence of CBX was not reported. Thus, we predicted that the movement of the Panx1 NTH out of the pore in PBN-blocked channels could create a favourable hydrophobic pathway for AEA (and CAY10455) to permeate the channel and that this would not occur in CBX pore-blocked channels.

We generated N-terminal deletion mutants by removing residues 2-20 of Panx1 (Panx1^ΔN20^)^19^ (**Fig. 4A**) from each ortholog (rPanx1^ΔN20^, mPanx1^ΔN20^, and hPanx1^ΔN20^) and expressed them in HEK 293T cells. Surface expression of ΔN20 was confirmed by biotinylation and was not significantly different from wild type (**Supplementary Fig 2**). Whole-cell patch clamp recordings revealed that the mutant Panx1^ΔN20^ had significantly smaller currents in response to voltage ramps compared to Panx1^WT^ (**Fig. 4B**), and no measurable CBX sensitivity (**Fig. 4C**), indicating that deletion of the NTH domain blocked ion conduction.

**Fig 4.**
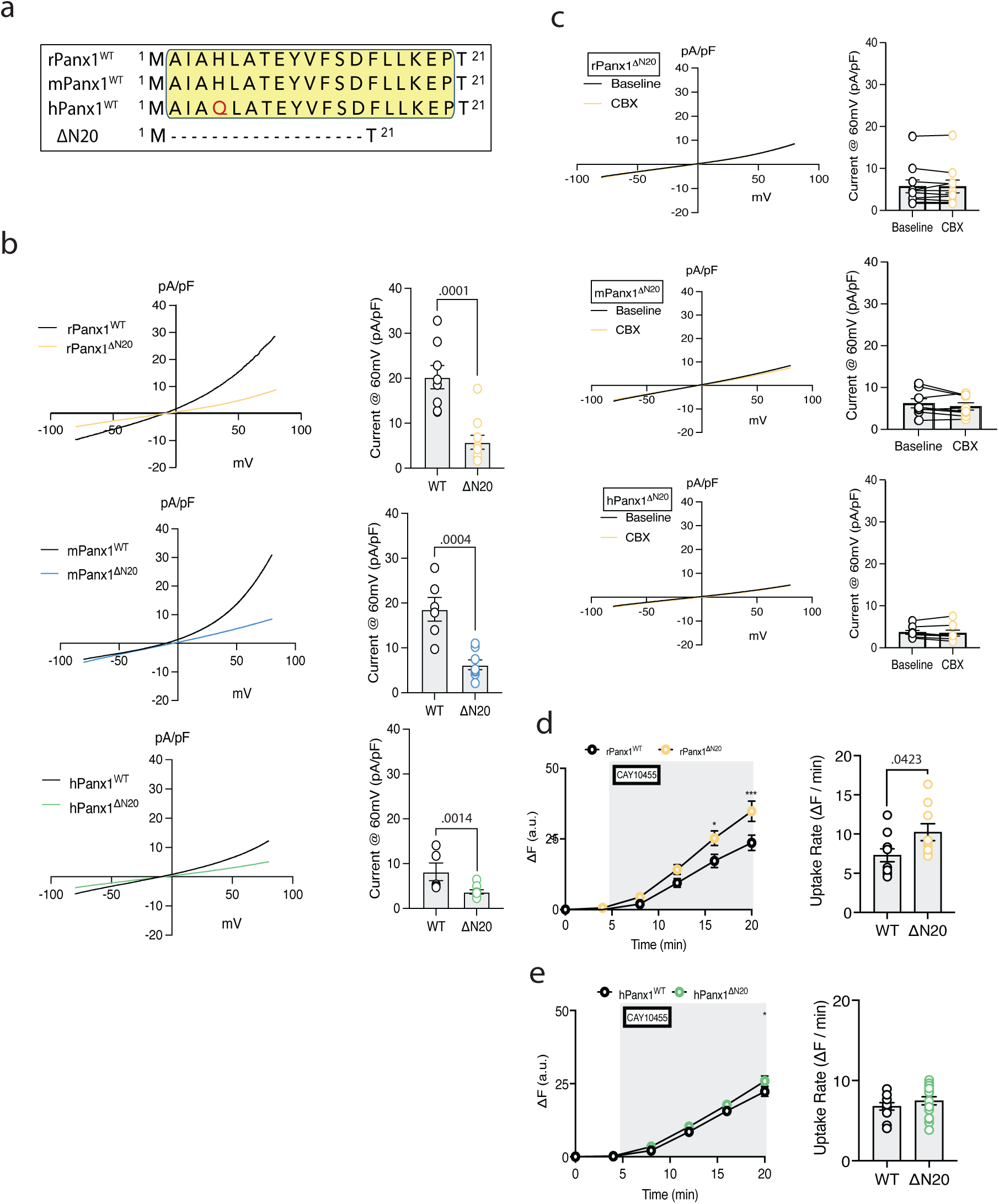
Deletion of the n-terminal helix of rPanx1 causes a loss of ion currents and an increase in CAY10455 uptake. **a**. sequence alignment of the rat mouse and human N-terminal domain of Panx1 and the region (2-20) of deleted amino acids for the ΔN20 mutant. **b**. comparison of whole cell currents for each wildtype (WT) and ΔN20 Panx1 ortholog. The left panels show exemplar whole-cell recordings, and the right panels compare current density (unpaired student’s t-test) for each ortholog and reveal significant inhibition of current with N-terminal deletion. (rPanx1^WT^ (n=8) vs rPanx1^ΔN20^ (n=10), P=0.0001, t=5.010; mPanx1^WT^ (n=6) vs mPanx1^ΔN20^ (n=8), P=0.0004, t=4.814); hPanx1^WT^ (n=8) vs hPanx1^ΔN20^ (n=5), P=0.0014, t=4.252). **c**. HEK 293T cells expressing the ΔN20 mutated Panx1 channels no longer show carbenoxolone (CBX; 100 μM) sensitive currents. Plots on the left are comparison of current density and no significant difference was observed between groups. (two-tailed paired student’s t-test. rPanx1 baseline vs CBX (n=10), P=0.8659, t=0.1737; mPanx1 baseline vs CBX (n=8), P=0.1519, t=1.608); hPanx1 baseline vs CBX (n=8), P=0.3798, t=0.9374). **d**. Left panel shows the time course of CAY10455 uptake by rPanx1 with and without the ΔN20 mutation. Experimental replicates were rPanx1 (n=10), rPanx1^ΔN20^ (n=9), (n=15). Right panels compares rates of uptake (two-tailed unpaired t-test (P=0.0423, t=2.195). **e**. Left panel shows the time course of CAY10455 uptake by hPanx1 with and without the ΔN20 mutation. Experimental replicates were hPanx1 (n=11) and hPanx1^ΔN20^ (n=15). Right panels compares rates of uptake (two-tailed unpaired t-test (P=0.3381, t=0.9774).

We next evaluated if the NTH deletion mutants had measurable CAY10455 uptake. As shown in **Fig. 4DE**, all three Panx1^ΔN20^ orthologs had substantive CAY10455 loading. Interestingly, rodent orthologs (rPanx1^ΔN20^ and mPanx1^ΔN20^) had increased CAY10455 uptake relative to WT constructs, whereas hPanx1’s uptake rate was comparable to hPanx1^WT^ (**Fig. 4D**). Overall, the deletion of the Panx1 NTH created a switch-of-function mutation that blocked ion conduction but maintained, or enhanced, CAY10455 uptake.

### Increased CAY10455 uptake by N-terminal deletion and probenecid are non-additive

As shown in Figs 3 and 4, PBN and NTH deletion increase CAY10455 uptake. Here we evaluated if PBN further enhanced CAY10455 uptake the rPanx1^ΔN20^ mutation. This was to evaluate if PBN and NTH deletion share a similar mechanism. Cells were exposed to PBN (1 mM) for 5 min prior to addition of 500nM CAY10455 in the continued presence of PBN. If PBN induces NTH movement out of the pore^19^, we predicted that PBN would not further increase CAY10455 in rPanx1^ΔN20^ expressing cells. PBN applied to rPanx1^ΔN20^ expressing HEK292T cells did not show increased CAY10455 uptake over that seen by NTH deletion alone (**Fig. 5AB**), suggesting they share a common mechanism.

**Fig 5.**
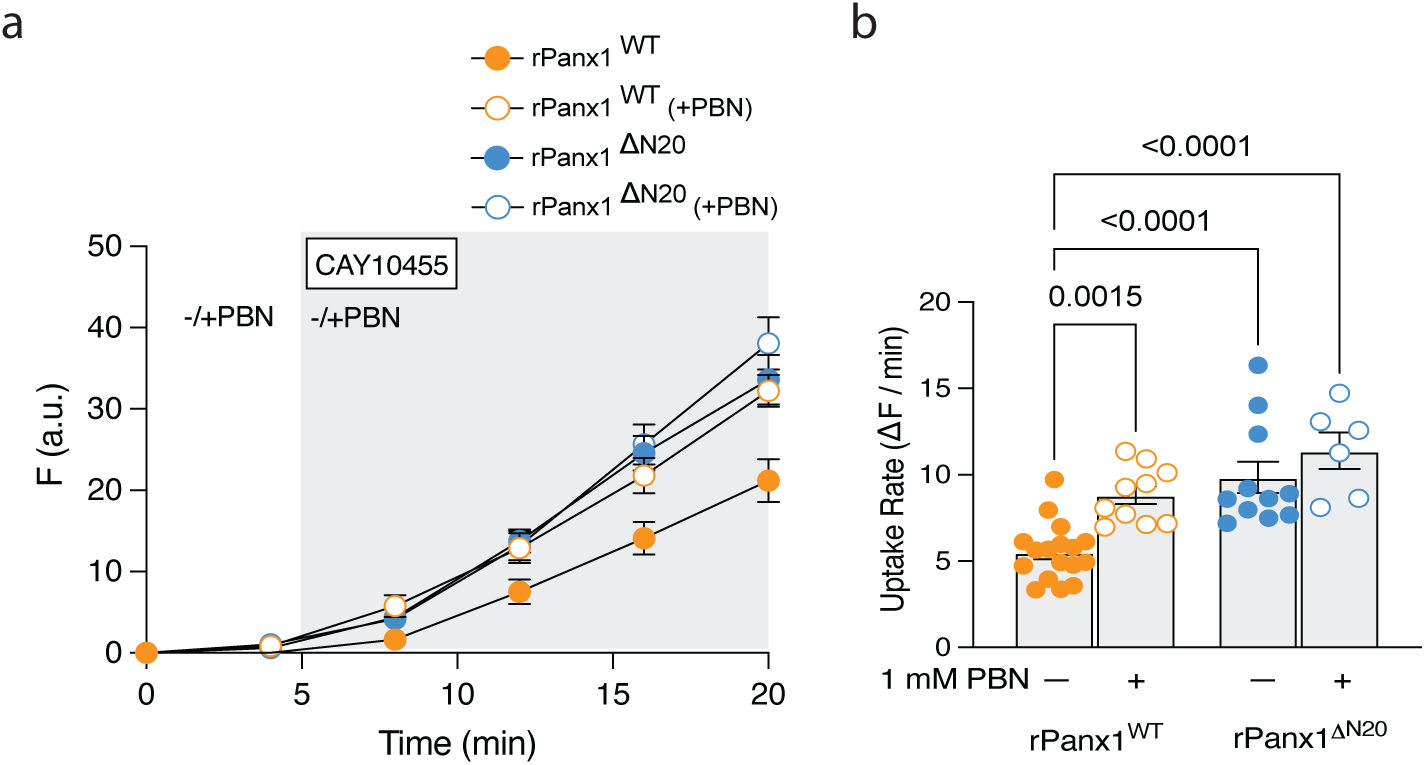
Probenecid block or N-terminal deletion of Panx1 share a common mechanism for the uptake of the AEA derivative, CAY10455. **a**. Time course of CAY10455 uptake by rPanx1, with and without the ΔN20 mutation in the presence or absence of probenecid (PBN; 1 mM). Experimental replicates are rPanx1^WT^ (n=17), rPanx1^WT^ +PBN (n=10), rPanx1^ΔN20^ (n=11), rPanx1^ΔN20^ +PBN (n=6). **b**. Rates of CAY10455 uptake calculated as the slope of the plots in a. Comparison between all groups was by one-way ANOVA (P<0.0001, F=19.70, df=56). Direct comparisons are indicated on the plot. Note that PBN and the ΔN20 mutation enhances CAY10455 uptake, and that the effect of PBN and ΔN20 are non-additive.

### Pore-lining I41 is important for ion conduction to AEA transport gating of Panx1

Multiple cryo-EM structures have been published showing lipid species embedded within the spaces between Panx1 subunits/homomers^18,19,28^. Specifically, Kuzuya et al resolved isoleucine residues (I41, I118, I278) that physically interacted with the lipids in inter-subunit spaces and suggested these amino acids are important in coordinating lipid movement into the Panx1 pore^19^. Here we sought to determine if I41, I118, and / or I278 were necessary for CAY10455 (and by extension AEA) transport by Panx1. Single, double, and triple point mutations of the hydrophobic isoleucine for polar serine (I41S, I118S, I278S, I118/278S, I41/118/278S) were generated with the prediction that loss of hydrophobicity would disrupt CAY10455 transport. All Panx1 orthologs have conserved isoleucine at these sites where I41 is pore facing and I118 and I278 are within the inter-subunit spaces (**Fig. 6A**). All the mutant rPanx1^(I118S, I278S, I118/278S, I41/118/278S)^ constructs had outward-rectifying CBX-sensitive ion currents (**Supplementary Fig. 3**), indicating that they were transported to the plasma membrane. Interestingly, the single point mutation rPanx1^I41S^ created a gain-of-function mutation that displayed a substantially larger CBX-sensitive current than that seen in the wild-type channels (**Fig 6B**). The other single mutants, double mutants and the triple mutant, rPanx1^I41/118/278S^ lacked this gain-of-function of ion conduction (**Fig. 6BC**). We tested all isoleucine mutant constructs in our CAY10455 uptake assay and found that contrary to the increased ion conductance, relative to rPanx1^WT^, CAY10455 uptake was significantly decreased in the rPanx1^I41S^, and rPanx1^I41/118/278S^ constructs (**Fig. 6DE**). Overall, this suggests that the rPanx1^I41S^ mutation independently creates a switch-of-function mutation with an opposite effect compared to the rPanx1^ΔN20^ mutation.

**Fig 6.**
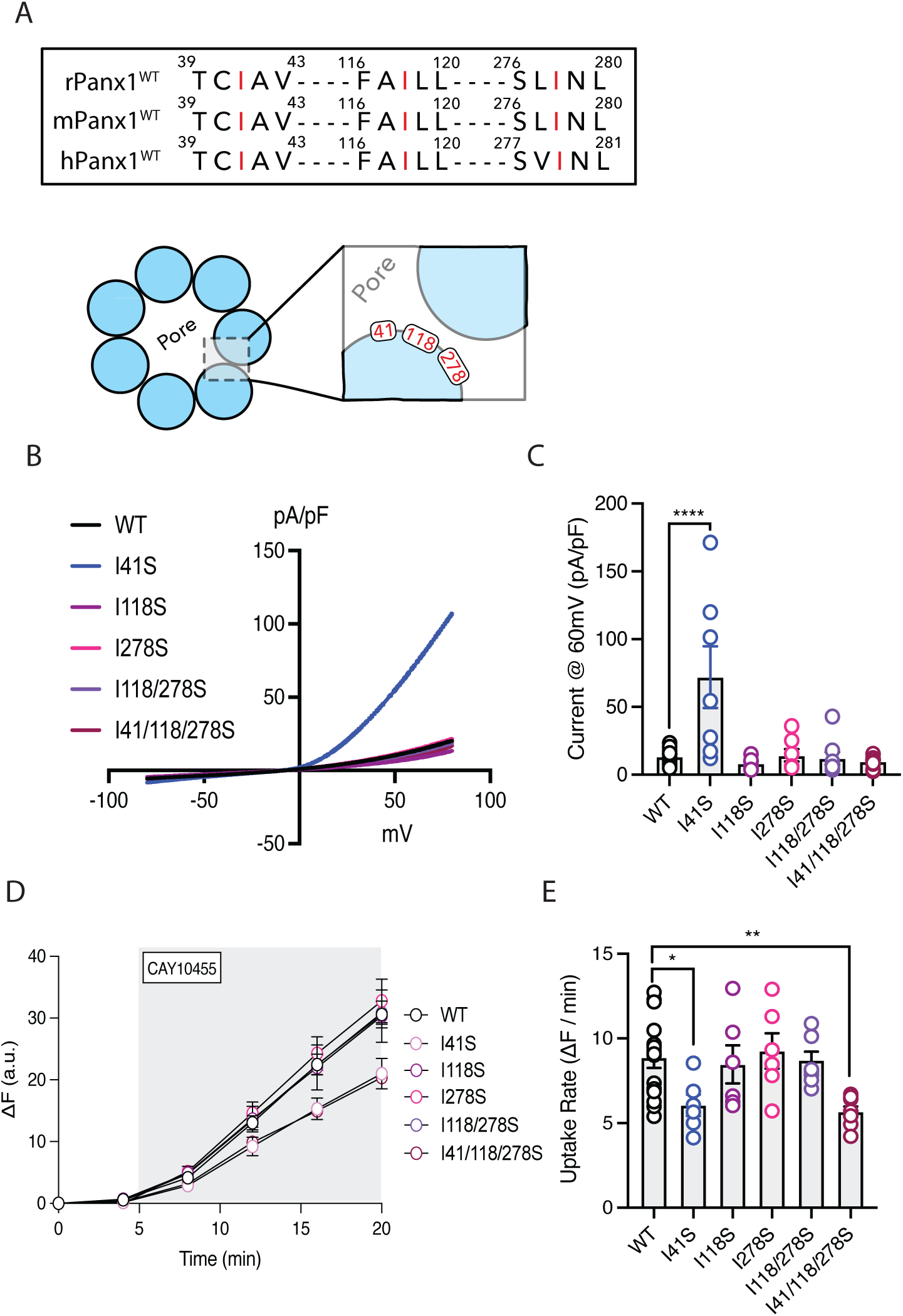
Mutation of the intrapore I41 residue creates a constitute Panx1 ion channel that lacks CAY10455 transport. **a**. Sequence alignment of the sites of isoleucine (red I) mutations to serine in the Panx1 orthologs and a schematic representation (lower) of their approximate location in the inter-subunit region of the heptameric channel. **b**. Whole cell recordings from HEK 293T cells expressing the single, double, and triple I to S mutations in rPanx1. **c**. Current density comparison of the I to S mutants. Experimental replicates are WT (n=10), I41S (n=7), I118S (n=7), I278S (n=7), I118/278S (n=8), I41/118/278S (n=10). One-way ANOVA indicates significant (P<0.0001, F=7.772, df=48) increases only in the I41S mutant. Direct comparisons are indicated in the plot. **d**. Time course of CAY10455 uptake by the I to S mutants. Experimental replicates are WT (n=15), I41S (n=6), I118S (n=6), I278S (n=6), I118/278S (n=8), I41/118/278S (n=8) **e**. Rates of uptake by the I to S mutants calculated as slope of the plot in d. Note the significant decrease in CAY10455 uptake in mutant channels carrying the I41S mutation. Comparison was by one-way ANOVA (F=4.404, P=0.0025, DF=48; n’s for each mutant: wildtype (WT) = 15, I41S=6, I118S = 6, I118/278S = 8, I41/I118/I278S = 8). Error bars represent SEM.

## Discussion

Here we used expression of Panx1 orthologs, mutagenesis, electrophysiology, and imaging of CAY10455 uptake by HEK 293T cells to investigate if Panx1 can transport AEA. We show that Panx1 has a unique function as a channel that conducts ions and transports AEA. To the best of our knowledge, this dual role, ion conduction and lipid-derived molecule transport, has not been reported for other channels. The switching of conductance appears to be dependent on the N-terminal helix of the channel, which is inserted into the pore during ion conduction and likely faces the cytosol during AEA transport. The data presented here support our previous suggestion that Panx1 functions as an AEA transporter in the CA1 region of the hippocampus^5^.

The identity of EMTs for AEA has been sought for several decades, with no definitive protein discovered to date^29,30^. In the context of synaptic AEA signalling, spatiotemporal regulation of its concentration is required to efficiently control brain network communication, and this implies the presence of a specific transport mechanism for clearing AEA. Our group has identified a novel role of Panx1 in controlling extracellular concentrations of AEA to modulate excitability of CA1 neurons in the hippocampus^5^. We demonstrated that when Panx1 was pharmacologically blocked or selectively knocked out from hippocampal pyramidal neurons, this elevated extracellular AEA and drove spontaneous glutamate release dependent on pre-synaptic TRPV1^5^. This indirectly suggested that Panx1 was permeable to AEA. Here, we have shown that uptake of a fluorescent AEA derivative, CAY10455 is enhanced by ectopic expression of three Panx1 orthologs, suggesting this is a conserved function amongst species.

AEA is uncharged, making it impossible to measure as current using conventional electrophysiological techniques. Therefore, we used HEK 293T cells, which express minimal hPanx1, and CAY10455 to quantify AEA transport across the plasma membrane. We showed that the over-expression of Panx1 orthologs increased the uptake of CAY10455 compared to non-transfected controls. It is important to note that in control cells, we observed basal CAY10455 uptake, suggesting two possibilities: 1) there are alternative mechanisms of uptake of AEA (i.e. passive diffusion through the membrane; unidentified proteinaceous transporters) or 2) the minor endogenous expression of hPanx1 in this cell line is responsible for uptake. While we favour the latter possibility, there is a reported diversity of AEA transport mechanisms among different cell types^31^ that could account for basal uptake.

Interestingly, all Panx1 orthologs, rPanx1, mPanx1 and hPanx1 showed similar rates of CAY10455 uptake, suggesting conservation of the AEA transport mechanism across species. The hPanx1 construct characteristically did not generate large voltage-sensitive currents^21,22^, despite having a clear AEA permeability. This indicates that hPanx1 has substantial AEA transporter activity compared to minor ion channel activity. This possibility warrants future investigation given that Panx1 studied in rodents has been reported to be important in neurodegeneration and synaptic plasticity and is permeable to ATP^6,7,32–34^.

It is important to note that our original suggestion that Panx1 is involved in AEA transport was challenged by Alhouayek et al, who claimed that in contrast to the hippocampus, Panx1 was not involved in AEA uptake in PC3 prostate cancer cells or SH-SY5Y neuroblastoma cells ^5,35^. However, the group based these findings on the observation that CBX did not inhibit [^3^H]-AEA uptake. This is consistent with our findings that CBX does not inhibit the uptake of AEA, and we demonstrate structural evidence for an alternative route of Panx1-dependent AEA uptake.

### Untagged AEA differentially modulates CAY10455 uptake

A potential caveat of using CAY10455 as a surrogate for AEA uptake is that they have different molecular weights (804.8 vs 347.5 Daltons, respectively). Thus, it was important to determine if unlabelled AEA competed with CAY10455 for transport. To test this, we used rPanx1^WT^ expressing 293T cells and co-applied increasing concentrations of AEA in a physiologically relevant range (1 nM, 10 nM, or 100 nM) in a constant concentration of CAY10455 (500nM). Nanomolar concentrations of AEA differentially affected CAY10455 uptake; low nanomolar levels (1 nM, 10 nM) increased CAY10455 uptake, whereas 100 nM AEA significantly inhibited uptake. This suggests that low nM concentrations of AEA may facilitate its own uptake, possibly by acting as a ligand on Panx1. Higher nM concentrations were required to compete with CAY10455 for transport and this supports the notion that Panx1 is transporting AEA. AEA could be interacting with the transmembrane regions of single Panx1 protomers, as suggested by cryo-EM structures of phospholipid binding sites to the heptameric channel^19^.

Panx1 theoretically has seven binding sites for lipids that could induce progressive conformational changes to increase AEA uptake. A similar progressive activation of ATP permeability upon c-terminal cleavage has been proposed by the Bayliss group using a hexameric Panx1 concatemer construct^36^.

Since 100 nM untagged AEA reduced CAY10455 uptake, we asked if this concentration impacted Panx1 ion currents to gain insight into the possibility that ions and AEA share part or all of the permeation pathway (i.e. pore). If AEA and atomic ions (i.e. chloride; Cl^-^) compete for pore access, we expected to see a reduction in current amplitude in the presence of AEA. When recording rPanx1^WT^ currents before/after perfusion of 100 nM untagged AEA, we observed no change in current density, which suggests that during voltage activation, atomic ions and AEA do not directly compete with one another for pore access.

### The Panx1 NTH position gates AEA uptake

Kuzuya et al unveiled a unique contribution of the Panx1 NTH to channel gating where its position in the channel pore promotes ion conduction and ‘flipping’ into the cytosol closes the channel^19^. They further suggested that when the NTH rearranged into the cytosol, lipids filled the pore, raising the intriguing possibility that this hydrophobic pore environment is permissive for AEA transport. It is important to note that although lipids have been described in the pore of several channels^37^, this has recently been challenged and may artificially increase lipid density in pores and decrease hydration^38^. Regardless, of whether lipid movement occurs (or not) into Panx1, we have presented several lines of evidence to support our model that rearrangement of the NTH is important for gating ion conduction and AEA transport: 1) Deletion of the NTH region (ΔN20) of Panx1 induced a loss of ion conduction and increased uptake of CAY10455. 2) PBN, which is reported to cause the NTH to exit the pore^19^, blocked ion conductance but CAY10455 uptake persisted. 3) Addition of PBN to the ΔN20 mutant Panx1 channels did not further increase CAY10455 uptake, suggesting a convergence of mechanism for AEA transport. 4) CBX, which blocked ion conductance but has not been reported to cause significant structural changes in Panx1 and likely acts a pore blocker^18^, did not affect CAY10455 uptake.

We propose that movement of the Panx1 NTH out of the pore creates an AEA permeable environment, that is impermeable to hydrated ions. Surprisingly, the rate of CAY10455 uptake robustly increased in rPanx1 and mPanx1 orthologs compared to hPanx1. The difference in rate among Panx1 orthologs could be tied to rPanx1 and mPanx1’s more pronounced ionic currents. When rPanx1 and mPanx1 cryo-EM structures become available, they may potentially shed light on subunit architecture around the NTH domain, which could explain these differences.

### Ion conducting gain-of-function mutation in Panx1 reduces AEA uptake

Three isoleucine residues embedded either within Panx1’s pore (I41) or within the inter-subunit spaces between protomers (I118, I278) bind lipids, suggesting they may be important sites for regulating AEA transport^19^. Isoleucine residues are typically not involved with protein function, although they can play a role in substrate recognition of hydrophobic ligands (i.e. lipids)^39,40^. Single (I41S, I118S, I278S), double (I118/278S), and triple (I41/118/278S) mutants of these residues were created, converting them into polar serine to change hydrophobicity. We demonstrated that these single, double, and triple mutant constructs have voltage-activated currents, which are CBX-sensitive. However, the pore-facing rPanx1^I41S^ mutation created a gain-of-function channel, increasing ion conductance relative to rPanx1^WT^.

The I to S constructs were evaluated for CAY10455 uptake and found that relative to rPanx1^WT^, each construct (single or triple mutant) where the I41S mutation was present, had a slower rate of CAY10455 uptake. This suggests that I41S is important for channel gating between ion conduction and AEA transport, possible through an interaction with the NTH. One possibility is that rPanx1^I41S^ mutation stabilizes the NTH in the pore with the consequence of increased ion conductance and decreased AEA transport.

### Putative pathways for AEA permeation

Our observations that the Panx1 pore blocker CBX does not alter CAY10455 uptake (but blocks ion currents) suggests AEA is not likely accessing the channel pore through the extracellular selectivity filter. This leaves 2 possibilities for how Panx1 is facilitating AEA uptake. The first possibility is lateral movement of AEA within the lipid bilayer and entry to Panx1’s pore between subunit interfaces. This would occur once the NTH has moved into the cytosol and allow AEA to bypass W74 in the extracellular facing selectivity filter. This movement of the NTH has been proposed to permit membrane phospholipids to migrate into the pore between Panx1 protomers and create a hydrophobic barrier to ion flux^19,27^ (see however ^38^). If this model is supported by future studies, it would suggest that there are structural elements of the inter-subunit interfaces that attract lipids to facilitate movement into the pore. It is possible that under physiological conditions, this mechanism is preferential for AEA over membrane lipids.

The second possible mechanism for AEA transport by Panx1 would not involve movement of AEA through the channel pore. Rather, it is possible that AEA interacts with lipid bilayer facing residues in the intramembrane space and is shuttled vertically through the membrane and into the cell. This model would be akin to lipid scramblases, such as TMEM16 family^41^. Once entering the cell through either pathway, it is likely that AEA then binds to fatty acid binding proteins or other carrier molecules to be delivered to FAAH for degradation^13^.

Overall, AEA is a lipid derived signalling molecule with activity across multiple systems in addition to the brain. Due to the variety of physiological roles for AEA, it follows that several uptake mechanisms may be present to meet the temporal and spatial requirements of each system. In this study, we define a putative mechanism where AEA uptake is regulated by the ion / metabolite channel Panx1. Our previous work has indicated that this uptake activity of Panx1 is important for regulating presynaptic glutamate release in the hippocampus by controlling AEA concentrations. Thus, the Panx1 mechanism could be restricted to systems that require rapid AEA clearance in small spaces such as synapses. However, given the almost ubiquitous distribution of Panx1 in tissues, it is conceivable that the channel plays a broader role in controlling AEA concentrations and its downstream signalling pathways.

## Methods

### Molecular Biology

*Rattus norvegicus* Pannexin-1 (rPanx1) and *Mus musculus* Pannexin-1 (mPanx1) complementary DNA (cDNA) was cloned between BamH1 and Sal1 sites in a pRK5 expression vector (pRK5-rPanx1, pRK5-mPanx1 respectively). *Homo sapiens* Pannexin-1 (hPanx1) cDNA was cloned between Nhe1 and Xho1 in a pcDNA3.1(-) expression vector (pcDNA3.1(-) – hPanx1).

Enhanced green fluorescent protein (EGFP) plasmid (pEGFP-C1) was commercially purchased (Addgene). The fluorescent protein mKate was inserted into a pcDNA3.1(-) expression vector (pcDNA3.1-mKate). Site-directed mutagenesis on cDNAs was performed using the Q5 site-directed mutagenesis kit (NEB) with the primers constructed using the NEBuilder tool (NEB). Primers used for site-directed mutagenesis constructs are found in Table.1. Mini-preparations of plasmid constructs were isolated using QIAprep Spin Miniprep Kit (Qiagen). Proceeding sequence confirmation, Maxi-preps of the plasmid construct were isolated using a Nucleobond Xtra Maxi Plus EF kit (Macherey-Nagel). The full coding frame of all pRK5-rPanx1 and pRK5-mPanx1 constructs were sequenced at the University of Calgary Sequencing Core using the primers CMV-F (5′-CGCAAATGGGCGGTAGGCGTG-3′) and pBABE3-R (5′-ACCCTAACTGACACACATTCC-3′) with annealing temps of 65°C and 56°C respectively. All pcDNA3.1(-)-hPanx1 constructs were sequenced using the primers CMV-F (5′-CGCAAATGGGCGGTAGGCGTG-3′) and pcDNA-R (5′-TAGAAGGCACAGTCGAGGC-3′) with annealing temps of 65°C and 58°C respectively.

### Cell Culture and Transfection

Human embryonic kidney-293T cells (HEK 293T) were purchased from ATCC and routinely maintained in the laboratory. Adherent HEK 293T cells were grown in Dulbecco’s modified Eagle’s medium + GlutaMAX (Thermo Fisher) supplemented with 10% fetal bovine serum (Thermo Fisher) and 1% penicillin-streptomycin (10,000 U/mL; Thermo Fisher) at 37°C in a 5% CO_2_ humidified growth incubator. HEK 293T cells between passages 5-20 were used for experiments. For patch-clamp recordings, HEK 293T cells reaching 40-60% confluency on 35mm culture plates were transiently transfected using Lipofectamine 2000 (Thermo Fisher) 36-48 hours before experiments, using the manufacturer’s protocol. HEK 293T cells were transfected with 2.5μg Panx1 and 0.5μg GFP cDNAs at a mixed ratio of 5:1 to identify positively transfected cells. For non-transfected cell controls, only 0.5µg GFP was used. For biotinylation experiments, HEK 293T cells reaching 40-60% confluency on poly-D-lysine (PDL)-coated 65mm cell culture dishes were transiently transfected using Lipofectamine 2000 36-48 hours before experiments, using the manufacturer’s protocol. HEK 293T cells were transfected with 2.5µg Panx1 cDNA construct. For non-transfected controls, only the transfection reagent was used. For fluorescent CAY10455 (Caymen Chemical) dye-uptake experiments, HEK 293T cells reaching 40-60% confluency on 35mm culture plates were transiently transfected using Lipofectamine 2000 36-48 hours before experiments. 293T cells were transfected with Panx1 (2.5µg) construct and 1µg mKate cDNAs to identify positively transfected cells. For a non-transfected control, only 1µg mKate was used.

### Cell-surface Biotinylation and Western Blots

Transfected HEK 293T cells overexpressing the Panx1 construct were washed twice in ice-cold HBSS (Ca^2+^/Mg^2+^ supplemented) prior to cell-surface protein labelling with EZ-Link Sulfo-NHS-SS-Biotin (Thermo). The biotin molecule was dissolved in ice-cold HBSS to a final concentration of 1 mg/mL and incubated on cells for 60 min at 4°C (kept away from light) before removing and quenching (192mM glycine, 25mM Tris, pH=8.3) the remaining biotin for 10min at 4°C (control cells/conditions were incubated with ice-cold HBSS only). Cells were washed twice in ice-cold HBSS and lysed in NP40 Lysis Buffer (Invitrogen) supplemented with 1X Halt Protease/Phosphatase Inhibitor Cocktail (Thermofisher) with end-over-end rotation at 4°C for 60 min. Lysates were centrifuged at 14000 rpm for 15 min with the soluble fraction quantified using the Micro BCA Protein Assay Kit (Thermofisher). 500µg protein lysate was volume normalized with NP40 buffer (inhibitor cocktail supplemented) and incubated with a 50% slurry of High-capacity Streptavidin Agarose Beads (Thermo) for 2 hours at 4°C with end-over-end rotation. Bead conjugates were centrifuged at 14000rpm and the soluble fraction was removed before performing three washes with NP40 Buffer and once with HBSS. Samples were eluted from beads at 95°C for 5min using 4X Laemmli Sample Buffer (Bio-Rad). Denatured samples from both the total lysate (20µg) and biotinylated surface fractions were separated on 10% Tris-HCl SDS-PAGE gels. Samples were transferred onto nitrocellulose membranes overnight at 4°C. Membranes were incubated in blocking buffer consisting of 5% bovine serum albumin (BSA; VWR) dissolved in 1X Tris-buffered saline (TBS) for 1 hour at room temperature prior to the addition of 0.05% Tween 20 (Bio-Rad) and primary antibodies overnight at 4°C. Primary antibodies included: 1:1000 anti-Transferrin-receptor (#13113S: Cell Signaling); 1:2000 anti-GAPDH (ab9484: Abcam); 1:2000 anti-Panx1 EL1 (Thompson lab produced); 1:1000 anti-MYC (#2278S: Cell Signaling); 1:1000 Phospho-Panx1^Y308^ (ABN1680 EMD Millipore; Thompson lab produced); 1:1000 Phospho-Src (Y416) (#6943S: Cell Signaling). Blots were washed 3 times in 1X TBS supplemented with 0.05% Tween 20 (TBST) prior to secondary antibody incubation.

Secondary antibody labelling was performed in 1X TBS containing 5% BSA, 0.05% Tween 20 and 1:10000 IRDye secondary antibodies (IRDye680 donkey anti-mouse and/or IRDye800 donkey anti-rabbit; LICOR) for 1 hour at room temperature. Blots were washed 2 times in 1X TBST, and once in 1X TBS prior to exposure on a LICOR Odyssey CLx imaging system.

### Patch-clamp Electrophysiology

On the day of experiments, transiently transfected HEK 293T cells overexpressing Panx1 construct and EGFP cDNAs were seeded onto poly-d-lysine coated glass coverslips at least 2 hours before recordings. HEK 293T cells were maintained at 30°C throughout the duration of each recording and voltage-clamped in whole-cell configuration using a Multiclamp 700B amplifier (Axon Instruments). Data was acquired using Clampex (v.10) software and an Axon Digidata 1550A digitizer (Axon Instruments) at 10 kHz, with currents analyzed offline in Clampfit software (v.10.7). Patch pipettes were pulled from 1.5/0.86mm [outer diameter/inner diameter] borosilicate glass (Sutter Instrument) using a P-1000 Micropipette Puller (Sutter Instrument) and had tip resistances of 3-5 MΩ. Patch pipettes contained (in mM) 4 NaCl, 1 MgCl_2_, 0.5 CaCl_2_, 30 TEA-Cl, 100 CsMeSO_4_ 10 EGTA, 10 HEPES, 3 ATP-Mg^2+^, and 0.3 GTP-Tris, pH=7.3 (with KOH), ∼290 mosmol/liter (solutes purchased from Sigma). Recordings were performed on a Zeiss Axio Observer Z1 inverted epifluorescence microscope using a 40x/0.6 air-immersion objective (Zeiss), 470nm light-emitting diode (LED) (Zeiss) and a 38 HE filter set (Zeiss) to visualize EGFP signal. HEK 293T cells were under continuous bath perfusion with extracellular solution containing (in mM) 140 NaCl, 3 KCl, 2 MgCl_2_, 2 CaCl_2_, 10 D-(+)-Glucose, and 10 HEPES, pH=7.3 (with NaOH), ∼305 mosmol/liter (solutes purchased from Sigma). Cells were voltage-clamped at −60 mV in whole-cell configuration and exposed to a 300ms voltage ramp (−80 to +80 mV) before stepping back to −60 mV to record Panx1 voltage-sensitive currents. Baseline currents were first measured in extracellular solution before perfusion of Panx1 blockers: 100µM carbenoxolone (CBX; sigma); 1mM probenecid (Invitrogen). In some experiments, 100nM, 1µM, 10µM, or 100µM AEA (Caymen Chemical) was perfused in normal extracellular solution into the recording chamber. Drugs and compounds were dissolved or diluted in extracellular solution. For ion replacement/selectivity experiments, patch pipettes contained the same internal solution. All patch recordings were adjusted for liquid junction potential where necessary (determined using Clampex).

### CAY10455 Imaging

On the day of experiments, transiently transfected HEK 293T cells overexpressing Panx1 and/or mKate cDNAs were seeded onto PDL-coated glass coverslips at least 2 hours before imaging. Cells were maintained at 30°C during imaging while under continuous bath perfusion with extracellular solution. Imaging was performed on a Zeiss Axio Observer Z1 inverted epifluorescence microscope using a 40x/0.6 air-immersion objective (Zeiss), 470nm LED (Zeiss) and a 38 HE filter set (Zeiss) to visualize CAY10455 fluorescence. mKate was visualized using a 590nm LED (Zeiss) and a 62 HE filter set (Zeiss). CAY10455 compound was diluted to 500nM in extracellular solution from a 1mg/mL stock for uptake experiments. Cells were imaged every 4 minutes during baseline conditions (absent of CAY10455) prior to perfusion of the CAY10455 compound for up to 20 min (dye in the bath for a total of 15 min). Panx1 blockers, CBX and PBN, were added during the 5min baseline and were continuously present during CAY10455 uptake. For experiments that included co-application of untagged AEA, CAY10455 was perfused simultaneously with AEA to assess competition for uptake. AEA was diluted to final concentrations of 1nM, 10nM, and 100nM in extracellular solution.

### Data analysis

For patch-clamp experiments, current amplitude was normalized to cell capacitance and reported as current density (pA/pF) when generating current-voltage (IV) plots. CAY10455 was quantified using Fiji in ImageJ to quantify fluorescence pixel intensity. Cells expressing mKate in the field of view were analyzed individually by creating an ROI and subsequently following them throughout the experiment. The baseline (no dye in the bath) and background fluorescent signals were subtracted at each time point and quantified as ΔF, which was equal to background subtracted difference in the fluorescence at time, t minus the initial fluorescence. CAY10455 rate of uptake (ΔF / min) was determined by fitting a linear curve to the pixel intensity values from the timepoints of 8min to 20min (first and last time point the dye intensity is captured). Fluorescence intensity changes for all individual cells in a field of view were averaged and this represents a single experimental replicate.

For biotinylation experiments, protein band densitometry images were constructed using ImageStudioLite and quantified using FIJI. Panx1 bands were normalized to membrane-localized Transferrin Receptor. Statistical comparisons and data figure generation was performed using GraphPad Prism 8 (GraphPad Software) where data presented are mean ± standard error (SEM). Data were compared with either student’s t-test or one way ANOVA and significance set a p<0.05 (*).

## Acknowledgments

This work was supported grants to RJT from the from Natural Sciences and Engineering Research Council of Canada, Canadian Institutes for Health Research, Heart and Stroke Foundation of Canada, and the Krembil Foundation.

## Author Contributions

CLA, ACN and AKJB performed all experiments and data analysis. CLA and FV designed primers for mutagenesis and proofed the final manuscript. CLA, NLW, ACN, AKJB and RJT designed the experiments and wrote the manuscript.

---

**Supplemantary Fig 1.**
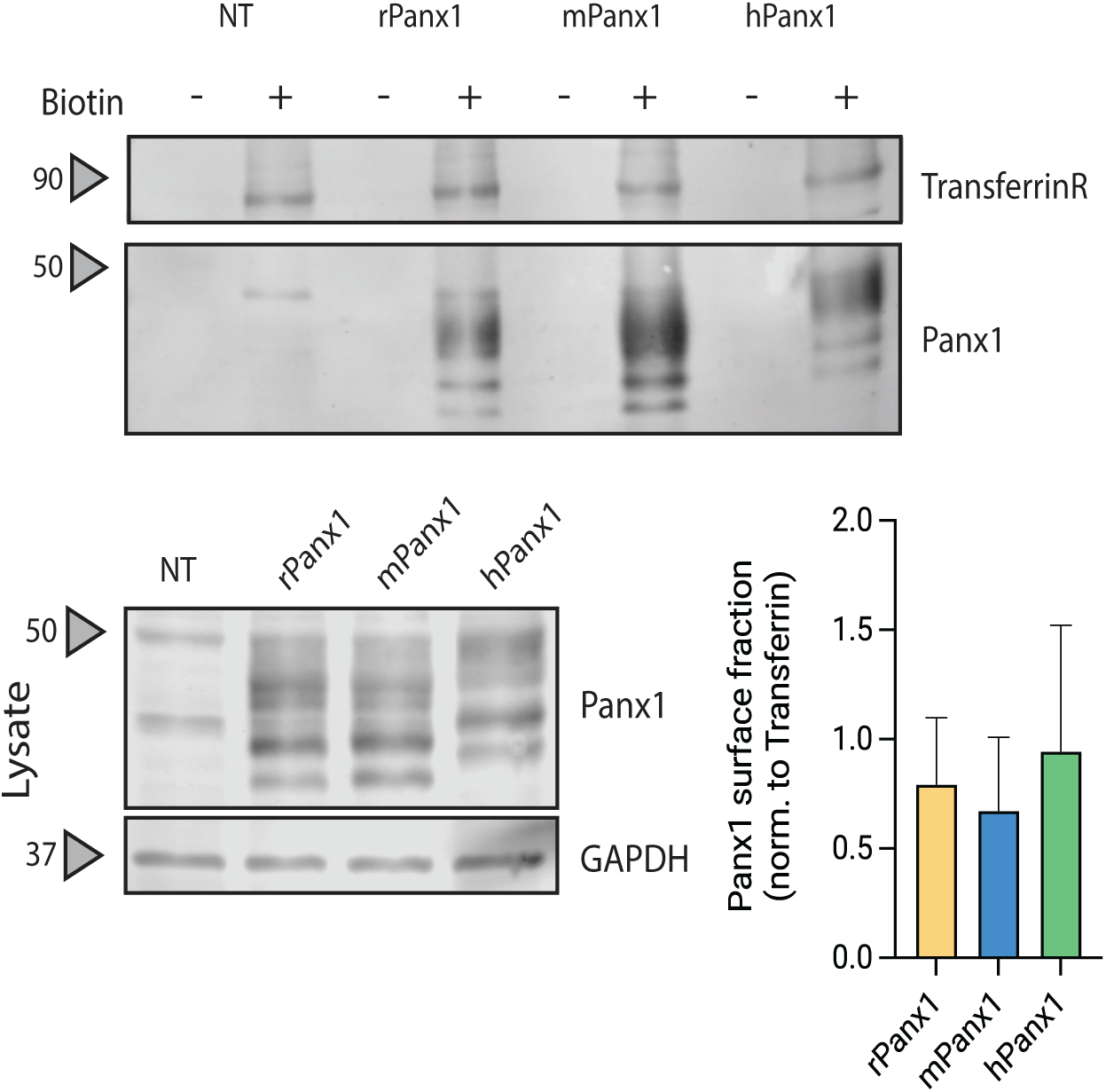
Cell surface biotynatlion of Panx1 orthologs expressed in HEK 293 T cells (top blot) compared to the whole cell lysate (bottom blot). Normalized expression of the 3 orthologs were not significnatly different. One-way ANOVA, P=0.9042, F=0.1042, df=5.

**Supplemental Fig 2.**
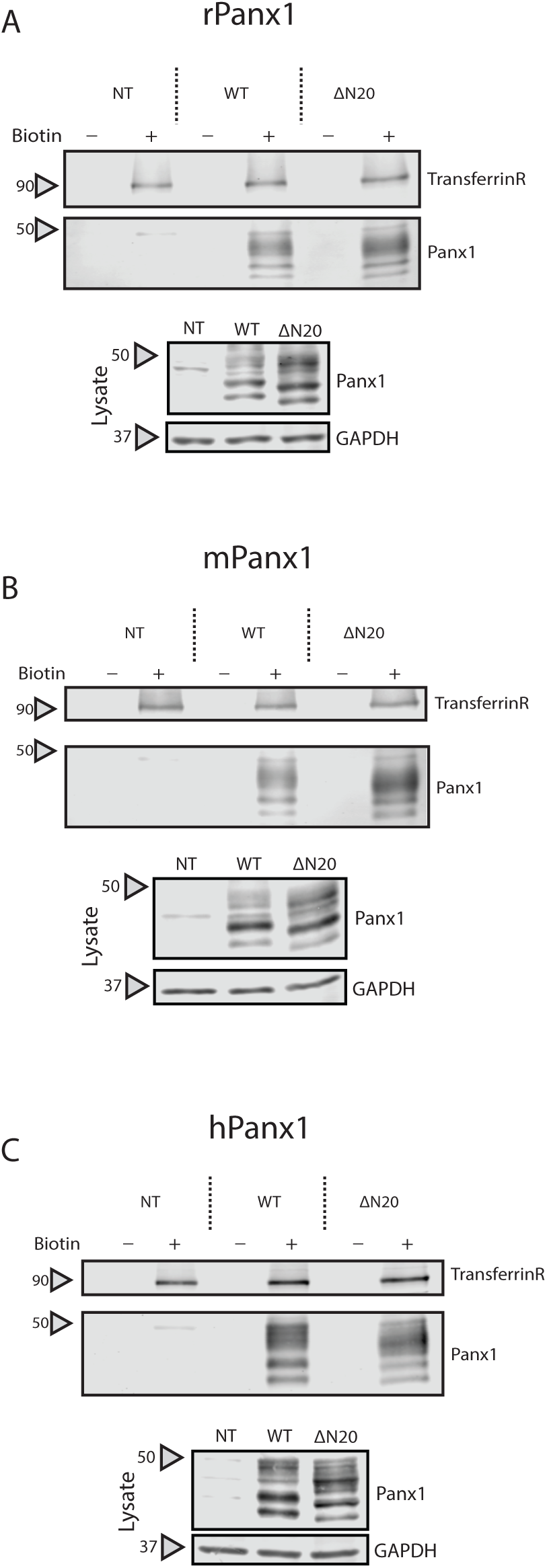
Example cell surface biotinylation Western blots showing that all 3 Panx1 orthologs and N-terminal helix deleltion mutants traffic to the cell surface.

**Supplementary Fig. 3.**
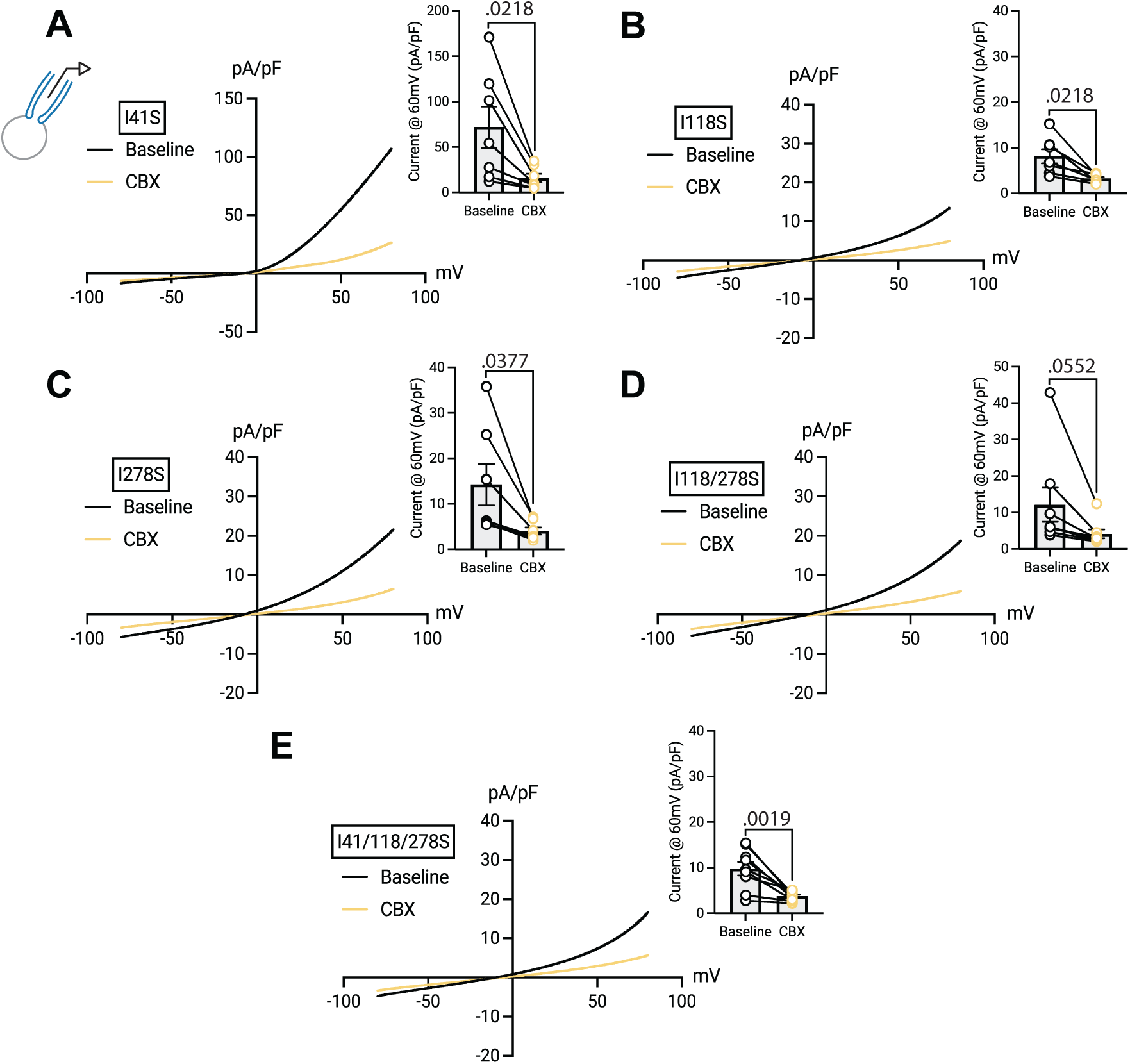
Whole cell, CBX-sensitive currents are observed with all single, double and triple isoleucine point mutations. All comparisons are by two-tailed paired t-test at +60mV between the baseline and with CBX (histogram insets). I41S n=7, P=0.0218, t=3.073. I118S n=7, P=0.0124, t=3.526. I278S n=7, P=0.0377, t=2.656. I118/278S n=8, P=0.0552, t=2.297. I41/118/278S n=9, P=0.0019, t=4.528.

